# Antibiotic Resistance Increases Evolvability and Maximizes Opportunities Across Fitness Landscapes

**DOI:** 10.1101/750729

**Authors:** Fabrizio Spagnolo, Daniel E. Dykhuizen

## Abstract

Antibiotic resistance continues to grow as a public health problem. One of the reasons for this continued growth is that resistance to antibiotics is strongly selected for in the presence of antibiotics and weakly selected against after their removal. This is frequently thought to be due to the effects of compensatory mutations. However, compensatory mutations are often not found in clinically relevant strains of antibiotic resistant pathogens. Here, we conduct experiments *in vitro* that highlight the role that fine scale differences in environment play in the maintenance of populations after selection for resistance. We show that differences in the mode of growth, dictated by environmental factors, are capable of reliably changing the force and direction of selection. Our results show that antibiotic resistance can increase evolvability in environments if conditions for selection exist, selecting differentially for newly arising variation and moving populations to previously unavailable adaptive peaks.

**Significance:** Antibiotic resistant bacteria are a large and growing problem for public health. A major question has been why antibiotic resistant strains do not disappear when they must compete with higher fitness drug sensitive strains. Here we show that selection for antibiotic resistant strains is particularly sensitive to differences in environmental conditions and that these differences help to define the fitness landscapes upon which these populations adapt. The result is an increase in evolvability, with many adaptive peaks that drug resistant populations can explore through natural selection, making predictions of evolution difficult and selection against resistant strains improbable.

## Introduction

One of the most difficult aspects of the antibiotic resistance problem is the contrast between the speed and strength of selection for antibiotic resistant strains and the slower selection against antibiotic strains of bacteria. The evolution for antibiotic resistance occurs either due to de novo mutations or due to horizontal gene transfer events. The concentration of antibiotics in the environment can either be low enough to allow the sensitive strains to survive but high enough to select for resistant strains or the concentration of antibiotics in the environment can be high enough that the drug susceptible phenotypes die off, leaving only resistant phenotypes to survive, grow, and reproduce. Different mutations are selected in these two cases (Wistrand-Yuen et al. 2018). In this paper we are studying the latter case, where only the resistant strains survive, because of high levels of antibiotics. Essentially, this is the strongest way that selection can act: either you are susceptible and quickly die, or you are resistant and survive. In laboratory experiments, the evolution of a population from susceptible to resistant has been shown to occur on the scale of hours to days (Spagnolo *et al*., 2016; Toprak *et al*., 2011; Miller *et al*., 2013).

As frequencies of antibiotic resistance rose worldwide, coordinated attempts were undertaken to reverse resistance by removing the environmental conditions that promoted it in the first place (Ridley et al. 1970). In essence, the strategy was to select against resistance as quickly and consistently as possible. This was done primarily by lowering rates of antibiotic usage and relying on the phenomenon known as the “cost of resistance” (Levin et al. 1997; Andersson and Levin 1999; Austin et al. 1999), whereby many resistant strains have lower fitness that the sensitive strains in the absence of antibiotics. Typically, this occurs because a vital phenotype, such as protein or membrane synthesis, is the target of the antibiotic, and therefore of the resistance mechanism as well. As such, the phenotype of resistant mutants is not like wild type, making them less fit in direct competition.

Results using this ecological approach have been mixed (Andersson and Hughes 2010), with reversion away from resistant phenotypes and back to antibiotic sensitivity shown to occur slowly (Seppala et al. 1997), quickly (van den Bogaard et al. 2000; Agersø and Aarestrup 2013), or not at all (Smith 1975; Mölstad et al. 2008).

Resistant mutations are reliably selected for when antibiotics are used for medical or agricultural purposes (Bush et al. 2011; The Review on Antimicrobial Resistance 2016), but do not always decline in frequency when the antibiotics are removed for the environment. It has been proposed that this is due to compensatory mutations (Maisnier-Patin and Andersson 2004). Compensatory mutations relieve the cost of resistance through partial restoration of the impacted phenotype (Reynolds 2000; Szamecz et al. 2014; Hughes and Andersson 2017) and have been consistently found in laboratory experiments (Schrag et al. 1997; Reynolds 2000; Maisnier-Patin and Andersson 2004). But, compensatory mutations are often absent from clinical strains of antibiotic resistant bacteria (Andersson and Levin 1999; MacLean and Vogwill 2015), suggesting that something more is at play.

In order better understand post-resistance adaptation in bacterial populations, we conducted several investigations using streptomycin resistant strains of *Escherichia coli* with a K42N mutation in ribosomal protein S12 under *in vitro* conditions with controlled differences between them. The results indicate that the adaptive landscape available to a population is of vital importance to how the population evolves, even when environmental differences seem to be of extremely fine scale, suggesting that how populations adapt in antibiotic resistance is even more intricate than previously feared.

## Methods

### Strains

All bacterial strains used are *E. coli*, K-12 MG1655 strains. The ancestral strain for all subsequent experiments is denoted as DD1953 and is known to be free of any plasmids, to be antibiotic sensitive and to be *rpoS-* due to a premature stop codon in the coding sequence of the *rpoS* gene. The DD1953 strain served as the streptomycin sensitive ancestral strain for all experiments. In order to assure genetic homogeneity at the start, DD1953 was grown from a single *E. coli* colony from an agar plate, grown in liquid media and then frozen for long term use.

We used a standard mutant screen to generate streptomycin resistant mutants of DD1953. The single nucleotide polymorphism (SNP) giving streptomycin resistance was identified and confirmed by MiSeq as well as Sanger DNA sequencing to be a nonsynonymous change in the *rpsL* gene at codon 42, changing the wild type leucine to asparagine (K42N). A pure freezer stock was prepared and stored at −80°C. This K42N *rpsL* mutant is the direct ancestor of all subsequent experimentally evolved populations. This resistant ancestor is labeled as FS1, or in the case where the mutant is also resistant to T5 bacteriophage, as FS5.

Resistance to the bacteriophage T5 was used as a molecular marker in situations where one was desirable, such as in direct competition experiments. Resistance to T5 is due to a SNP in the *fhu* gene and has been shown to be selectively neutral in chemostat experiments (Moser 1958). The minimum inhibitory concentrations (MIC) for all strains were found experimentally using methods identical to those described earlier (Spagnolo et al. 2016). We found that DD1953 had a MIC for streptomycin of 2-4 *µ*g/mL, with all resistant strains having MICs over 1000 *µ*g/mL.

### Chemostats and Serial Transfer Flasks

To inoculate the chemostats and flasks, we grew strains FS1 and FS5 overnight in flasks inoculated from frozen cultures containing Davis salts and 0.1% glucose at 37°C shaken at 200 rpm. These cultures were diluted the next morning into fresh media and the growth followed by measuring optical density. When the optical density reached 1.7, we added 1ml of each inoculating strain to chemostats and flasks containing 30ml of experimental media, which was Davis salts supplemented with 0.01% glucose (w/v) as the only available carbon source. Streptomycin concentrations for all experimental evolution vessels were as indicated.

All continuous culture experiments were conducted in Kubitschek chemostats following procedures previously described (Dykhuizen 1993). Here, the generation time was set to 3.69 hours per generation (6.5 gen/day, D = 0.1875), matching the mean number of generations per day in the 1:100 dilution of flask cultures. Additional chemostat related methods are in the Supplemental Information (**SI**). A total of 4 chemostats were run simultaneously. Chemostats 1 & 2 utilized the Davis minimal media with streptomycin at a concentration of 100 μg/mL (high streptomycin condition). Chemostat 3 had the Davis minimal media and 16 μg/mL of streptomycin (low streptomycin condition) as per Miller (Miller 1992). Chemostat 4 had Davis minimal media but contained no streptomycin (no streptomycin condition). All streptomycin came from a single prepared stock (10 mg/mL).

Two serial transfer flask experiments were run concurrently with the chemostat experiments. Whereas the chemostats maintained the experimental populations in a constant state of sub-maximal exponential growth, the serial transfer flask populations underwent full cycles of lag phase, maximal exponential growth, and stationary phase each day. Two sterile 250 mL flasks were prepared with 30 mLs of fresh Davis minimal media, 0.01% glucose, and 100 μg/mL of streptomycin (high streptomycin condition). The flasks were placed in an incubator, maintained at 37°C and constantly shaken at 200 rpm for 24 hours. After a 24 hour period, 300 μL of the experimental sample was transferred into a fresh 250 mL flask with 30 mL of fresh media. The 100 fold dilution regime and 24 hour time frame allows for the calculated number of generations to be 6.5 generations, as per Lenski (Lenski et al. 1991). This experimental design allows the *average* generation time of the *E. coli* in the flasks to match the *maintained* generation time in the chemostats.

### MiSeq

Miseq NG sequencing was performed on population samples. Libraries were prepared using the Illumina TruSeq genomic DNA-nano and sequenced using Illumina MiSeq 2×300 protocols at the University of Wisconsin Biotechnology Center (Madison, WI). Overall mean coverage of the genomic sequencing was 162X (SD: 28).

### Bioinformatics

All bioinformatics was performed in the Galaxy environment. The raw sequence reads were vetted for quality using FastQC and aligned to the ancestral genome (DD1953) using the reference K-12 MG1655 genome as a scaffold (Genbank accession NC_000913.3). SNP calling and indel identification were conducted via the VarCap workflow (Zojer et al. 2017), which also provides SNP frequency data.

### Protein Synthesis Assay

Streptomycin acts by binding near the A-site of the ribosome, interfering with the function of protein S12, the product of the *rpsL* gene. The mechanism of action for streptomycin is interfering with protein synthesis efficiency, both in rate of protein synthesis and fidelity (Kurland 1992). Structural changes to the S12 protein, particularly at amino acid position 42, are known to result in high level resistance to streptomycin, but result in reduced protein synthesis efficacy.

In order to compare protein synthesis rates for streptomycin sensitive, resistant, and evolved strains, we performed a protein synthesis assay. S30 cell lysate was extracted from an exponentially growing population of *E. coli* that was to be assayed. The S30 lysate extraction protocol was based upon that of Shrestha (Shrestha et al. 2012) with some modification (See SI). Following S30 lysate extraction, a firefly luciferase assay was performed as per the *E. coli* S30 Extract System for Circular DNA by Promega guidelines for control experiments using the Promega pBEST*luc* plasmid for *E. coli* (Promega, Madison, WI). Following 60 minutes of incubation at 37°C, relative light units (RLU) were measured in triplicate using a Turner Biosystems (Sunnyvale, CA) Model 20/20n luminometer (See SI).

### Fitness Assay

In order to compare relative fitness of resistant and evolved strains, direct competitions between strains were conducted in chemostats in the manner of Dykhuizen (Dykhuizen and Hartl 1980; Dykhuizen 1993). Using the T5 bacteriophage molecular marker, two strains were inoculated in a chemostat in equal numbers. Changes in the frequencies of these strains were tracked over 72-96 hours via plating as described.

By plotting the natural log of the ratio of mean CFUs per unit time, the coefficient of selection was obtained from the slope of the plotted trendline. From these measures, the relative fitness measure, w_R_, was calculated by use of the equation:

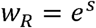

All chemostat parameters were as described but the concentration of streptomycin was varied to coincide with the context of the competition. We note that competition was under chemostat conditions even for strains evolved in flasks.

### Growth Curve Assay

We obtained growth curve data for strains by monitoring optical density (λ = 600 nm) in 96-well plates in a SpectraMax Model 384 Plus microplate reader using SoftMaxPro software (Molecular Devices, Sunnyvale, CA) over a ten hour period. Briefly, we grew strains overnight in 3 mL of LB (rich media). The next day, 300 μL of the overnight growth was added to 9 mL of Davis minimal media (1:30 dilution). The media-strain mixture was vortexed to insure even distribution. The inoculated media was then put into replicate wells on the 96 well plate, with 180 μLs in each well and at least 1 negative control well per set of technical replicates. A minimum of 16 replicates were run for each sample.

Following inoculation, the 96 well plate was placed into the plate reader and the populations were allowed to grow. The OD was measured every 60 seconds over a period of 10 hours, with mixing of the plate between reads. Typically, the populations reached stationary phase well before the 10 hour mark. Data was analyzed using the grofit package (Kahm et al. 2010) and R statistical software (R Development Core Team 2012). The grofit package determines the time point for transition into exponential growth as well as into stationary phase, thereby allowing for analysis of length of time spent in lag and exponential growth phases, as well as the maximum growth rate (μ_max_), and the density at the transition into stationary phase.

The amount of time spent in exponential growth was calculated by finding the difference between the time point where a well population entered stationary phase and the time point where exponential phase was entered.

## Results

Based upon the results of our fitness assays as well as the substantial body of previous work regarding mechanisms of fitness increases, particularly with regard to streptomycin resistance, we expected our experimental results would confirm previous work. This presumption was only partially correct.

### Loss of Fitness in Streptomycin Resistant Mutants

Amino acid changes at position 42 of the ribosomal protein S12 confer resistance to high doses of streptomycin. This mutation also confers substantial fitness costs in *Escherichia coli* (Kurland 1992), *Mycobacterium tuberculosis* (Sander et al. 2002) and *Salmonella enterica* (Bjorkman et al. 1998). When two *E. coli* strains with spontaneous streptomycin resistant mutations (both mutations were in codon 42 in the *rpsL* gene) were individually competed against an isogenic susceptible strain in flask cultures of a modified Davis salts medium with limited glucose (Schrag and Perrot 1996; Schrag et al. 1997), the selection found was 11 to 19% against the resistant strain.

In addition, Paulander *et al*., using a streptomycin resistant mutation in *Salmonella typhimurium* LT2 [K42N in the S12], measured maximal growth rate in pure culture and then compared growth rates of the strep-resistant strain to the sensitive strain to determine relative fitness (Paulander et al. 2009). The sensitive strain grew faster on a rich media (LB) and on M9 minimal media supplemented with glucose or with glycerol, but grew more slowly on poor media—M9 MM supplemented with pyruvate or with succinate. They showed this was due to the differential induction of the stress-inducible sigma factor, RpoS. On the poor media the strep-sensitive cells induce *rpoS*, which retards growth. When both cells lack *rpoS*, the effect disappears. This example shows how the effects of the environment and other physiological systems in the cell can reverse expectations.

We competed streptomycin resistant mutants, FS1 and FS5, (Table 1) in glucose limited chemostats. These mutations were in the codon 42 of the S12 ribosomal protein like the above example. Strains FS1 and FS5 had lower relative fitness when compared to the streptomycin sensitive ancestor, DD1953. The fitness of FS1 (T5 bacteriophage susceptible) and FS5 (T5 bacteriophage resistant) was measured to be 0.91 ±SD 0.02 relative to DD1953 (mean of three samples) in our fitness assay (Table 3). This level of fitness loss is in line with that found in previous experiments (Schrag and Perrot 1996; Björkman et al. 2000).

**Table 1:**
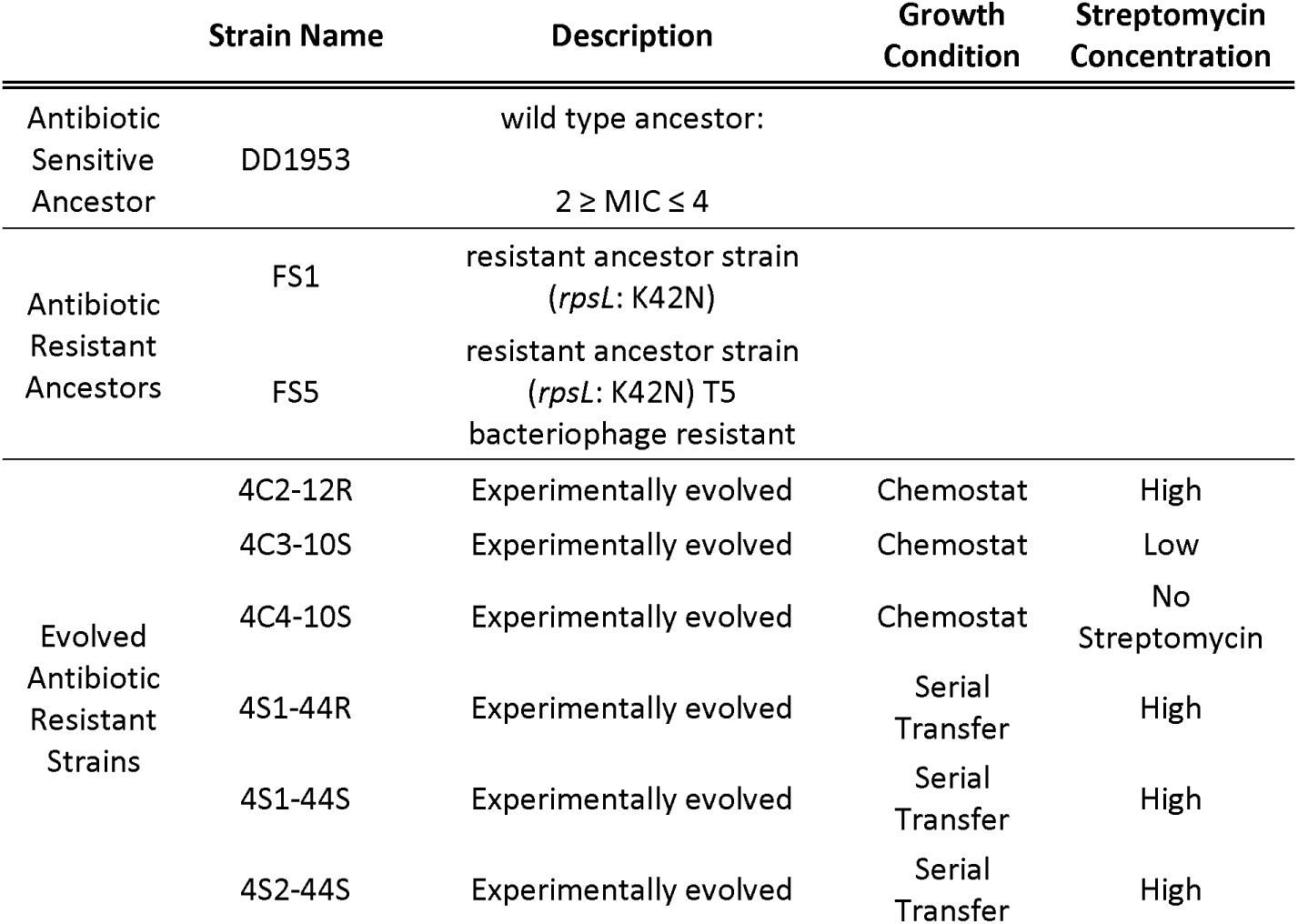
Strain Names and Experimental Conditions. All strains were derived from a single antibiotic sensitive ancestor, DD1953. Two streptomycin resistant mutants of DD1953 were generated, FS1 and FS5, differing only in resistance to T5 bacteriophage, which was used as a neutral molecular marker. Evolved strains experienced different *in vitro* environments (4C designates chemostat evolved and 4S means evolved in serial batch culture). Chemostat evolved strains also had a range of set streptomycin concentrations for their time under experimental conditions (last column).

**Table 2:** Genetic Changes in Evolved Strains Relative to Antibiotic Resistant Ancestor. Genetic changes identified through MiSeq relative to the relevant antibiotic resistant ancestor (FS1 or FS5). SNP frequency, as quantified through the VarCap bioinformatics pipeline, as well as function data (as per the Ecogene database, http://ecogene.org) are provided for each mutated gene.

| Evolved Strain | Gene and Change | Frequency | Description |
| --- | --- | --- | --- |
| 4C2-12R | *galS* Deletion | 0.14 | 1,022 bp deletion |
|  | *galS* Deletion | 0.26 | 1 bp deletion |
|  | *malT* | 0.41 | Nonsynonymous change |
|  | *malK* | 0.43 | Nonsynonymous change, ATP-binding subunit |
| 4C3-10S | *galS* | 1 | Nonsynonymous change |
| 4C4-10S | *galS* | 1 | Synonymous change |
| 4S1-44R | *rpsD* | 0.27 | Structural constituent of ribosome; nonsynonymous change, L198P. Previously unreported compensatory mutation. |
|  | *rpsD* | 0.2 | Structural constituent of ribosome; nonsynonymous change, I199S. Previously unreported compensatory mutation. |
|  | *rpsE* | 0.5 | Structural constituent of ribosome; nonsynonymous change, G108V. Known compensatory mutation site. |
|  | *acs* | 0.6 | Acetyl CoA synthase, acetate-scavenging enzyme, improves carbon starvation survival; promoter region |
| 4S1-44S | *rpsE* | 1 | Structural constituent of ribosome; nonsynonymous change, G108A. Known compensatory mutation site. |
|  | *yjcO* intergenic region | 0.27 | Sel1 family TPR-like repeat protein; function unknown |
| 4S2-44S | *gstB* promoter region | 1 | glutathione-dependent bromoacetate dehalogenase |
|  | *rpsE* | 1 | Structural constituent of ribosome; nonsynonymous change, G86C. Known compensatory mutation. |
|  | *yjcO* intergenic region | 0.26 | Sel1 family TPR-like repeat protein; function unknown |
|  | *yjeJ* proximal promoter region | 1 | Uncharacterized protein |

**Table 3:** Strain Relative Fitnesses before and After Evolution Experiment. All relative fitnesses were quantified from direct competition in chemostats. Fitness was measured as the slope of the best fit line for a time series of ratios of T5 sensitive to T5 resistant colonies (in manner of Dykhuizen and Hartl, 1980). A. Fitnesses of streptomycin resistant ancestors (FS1, FS5) relative to their streptomycin sensitive ancestor, DD1953. B. Relative fitnesses of all evolved strains relative to their pre-experiment ancestor, as obtained by direct competition. ND indicates not determined.

| Pre-Experiment |  | Post-Experiment |  |
| --- | --- | --- | --- |
| Strain | Fitness | Strain | Fitness |
| DD1953 | 1 | FS1 | 1 |
| FS1 | 0.919 | FS5 | 0.998 |
| FS5 | 0.917 | 4C2-12R | 1.04 |
|  |  | 4C3-10S | 1.03 |
|  |  | 4C4-10S | 1.03 |
|  |  | 4S1-44S | 1.02 |
|  |  | 4S1-44R | 1.02 |
|  |  | 4S2-44S | ND |

### Fitness Increases via Compensatory Mutations in Batch Culture

As in previous experiments with streptomycin resistant bacteria in batch culture, the streptomycin resistant mutants rapidly selected for compensatory mutations in the *rpsD* and *rpsE* genes (S4 and S5 small ribosomal proteins, respectively) (Table 2) and increased in fitness relative to the ancestor (2-4%). The compensatory mutations did not affect MICs of our strep-resistant strains.

Ribosomal ambiguity mutations (ram) in *rpsD* and *rpsE* have been shown to be compensatory (Maisnier-Patin et al. 2002) for hyperaccurate *rpsL* mutants, such as ours. This is believed to be because of ram mutants’ pleiotropic effect of increasing polypeptide synthesis rates, thereby counterbalancing the increased synthesis time caused by slowed proofreading in hyperaccurate mutants of *rpsL* (Bohman et al. 1984) (See Poehlsgaard and Douthwaite 2005; Holberger and Hayes 2009 for reviews). These ram mutations do not increase polypeptide synthesis when the genetic background includes a wild type *rpsL* gene, as measured *in vitro* using radioactive isotope tagged amino acids (Andersson et al. 1982).

There can be two approaches to testing the overall fitness effects of these compensatory mutations. The first was taken by Schrag and Perrot where they used P1 to transduce both the streptomycin sensitive and the streptomycin resistant alleles into the evolved cultures carrying the streptomycin resistant mutation (Schrag and Perrot 1996). Their results were quite variable but in all cases the strain carrying the resistant allele had an advantage over the strain carrying the sensitive allele in flask cultures. They found compensatory mutations in *rpsD* and *rpsE* which were selected in conjunction with a change at position 42 in the S12 ribosomal protein, but were detrimental when paired with the streptomycin sensitive allele. These compensatory mutations led to a condition where reversion to sensitivity would be selected against; genetic background matters in determining the fitness of the strain carrying a mutation giving resistence to streptomycin.

We took the second approach. The evolved strains carrying the mutation giving streptomycin resistance were competed against unevolved strains carrying the streptomycin resistance allele. The evolved strains had all been evolved either in flask culture or in chemostats (Table 1). Overall, evolved strains increased fitness relative to their streptomycin resistant ancestors 2-4% above where they started in the experiments (Table 3).

### No Previously Identified Compensatory Mutations in Chemostats

In our chemostat experiments, the growth medium was identical to that of the flask experiments, although the growth conditions clearly were not. We also varied the concentration of streptomycin in different chemostats to test for effect. Fitnesses also increased in these evolved strains, but the genetic mechanism was different. In chemostat cultures, we found no compensatory mutations in *rpsL, rpsE*, or *rpsD* (Table 2), all of which have previously been associated with changes in *rpsL*. By performing whole-genome sequencing of evolved clones, we identified changes in the glucose uptake systems of the chemostat evolved strains, particularly in *galS*, which have previously been found in chemostat evolved strains of strep-sensitive *E. coli* (Notley-McRobb and Ferenci 1999). Thus, the changes were not compensatory.

### Unexpected Phenotypic Changes

We performed growth curve assays in order to better understand the nature of the observed fitness increases. The expectation was that we would observe increases in maximum growth rates in the flask-evolved strains as a result of the compensatory mutations increasing protein synthesis efficiency. However, even prior to the experimental evolution experiments, maximum growth rates did not differ between strep-sensitive DD1953 and the resistant clones FS1 and FS5 in minimal glucose media. In the evolved strains, we were surprised to find no discernible pattern of increase in maximal growth rates, regardless of whether strains evolved in flasks or chemostats (Fig 1B). We note that the low maximum growth rate of strain 4C2-12R, which could be because of some unknown mutational or regulatory change, seems to be reflected in longer time in exponential growth. This could be a unique phenomenon for which we have no explanation. Additionally, there was no observable pattern to changes in time spent in lag phase or the population density at entry to stationary phase for evolved strains when compared to the strep resistant ancestors (Fig 1A&C).

**Figure 1:**
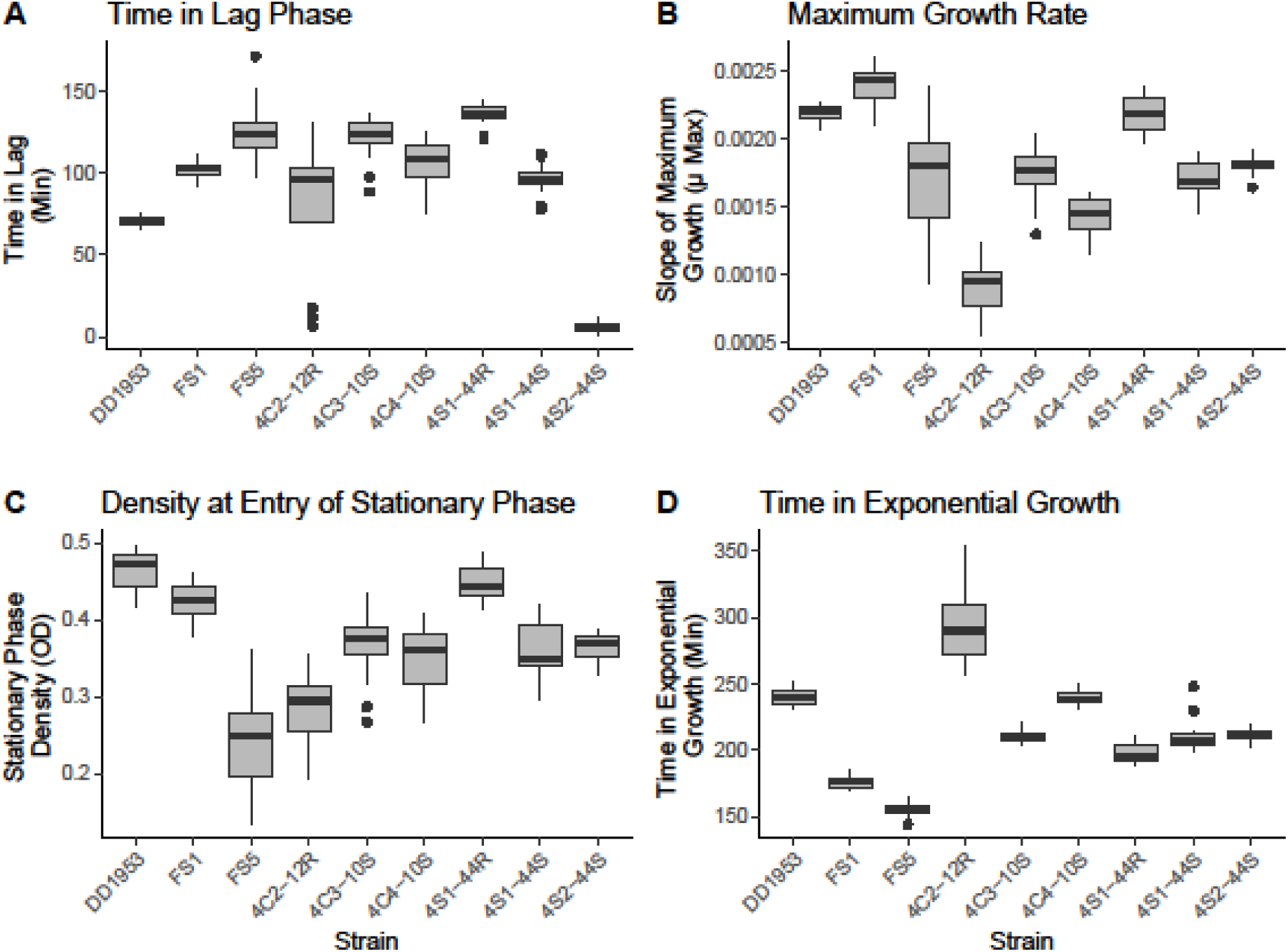
Boxplots of Growth Curve Assay for Ancestor and Evolved Strains. All strains were tracked in a 96-well plate assay for growth related phenotypes. A. Time Spent in Lag Phase: No observable pattern for changes in time spent in lag phase was found for strains. B. Maximum Growth Rate: Counter to expectation, maximal growth rate did not decline for streptomycin resistant strains FS1 or FS5 relative to DD1953 and then did not increase after evolution under serial passage conditions. In fact, *µ*_max_ declined in flask evolved strains relative to the antibiotic resistant ancestors. C. Population Density at Entry to Stationary Phase: The final density of experimental populations did not show a pattern after experimental evolution. D. Time Spent in Exponential Phase: The time spent in exponential growth significantly declined for streptomycin resistant mutants FS1 and FS5 relative to the DD1953 Ancestor. Following *in vitro* evolution, all evolved strains showed significant increases in the time spent in exponential growth, with values approaching those of DD1953. This phenotype has not been previously described relating to antibiotic resistance.

We did, however, find strong support for directional selection on the amount of time spent in exponential growth for all evolved strains. This increase in the time spent in exponential growth is consistent across all strains regardless of *in vitro* method, the concentration of streptomycin present, or the type or number of mutations found in the evolved populations. Our results indicate that regardless of experimental condition, selection consistently acts upon the time in exponential growth. FS1 and FS5, the unevolved strains containing new mutations giving rise to streptomycin resistance, have a normal growth rate but a shortened time in exponential growth. In the evolved strains derived from these two strains the time in exponential growth is like the original strain, DD1953, as if selection acts upon this phenotype.

In order to better understand these results, a functional protein synthesis assay was conducted. This assay also yielded unexpected results. Rather than indicating an increase in the speed and/or accuracy in producing functional proteins (particularly after compensatory mutations), the rate of synthesis of functional proteins for all evolved strains was *lower* than the streptomycin resistant ancestors in the presence of streptomycin (Fig 3). This indicates that the rate of protein synthesis, long thought to be the phenotype most effected by streptomycin resistance, may not tell the entire story of streptomycin-resistance adaptation in bacteria. As a control, we also performed the protein synthesis assay under strep free conditions. Under these conditions, DD1953, the strep-sensitive ancestor, produced functional protein at a rate 2,372X higher than when in the presence of streptomycin (Fig S1). For evolved strains, most showed little or no significant difference to results with streptomycin present. Two of three flask evolved strains had increases in functional protein synthesis when streptomycin was absent, but this is an environmental condition in which they did not evolve and the rate of protein synthesis was still approximately two orders of magnitude lower than in wild type. Additional data for the protein synthesis assays is in the Supplemental Information.

**Figure 3:**
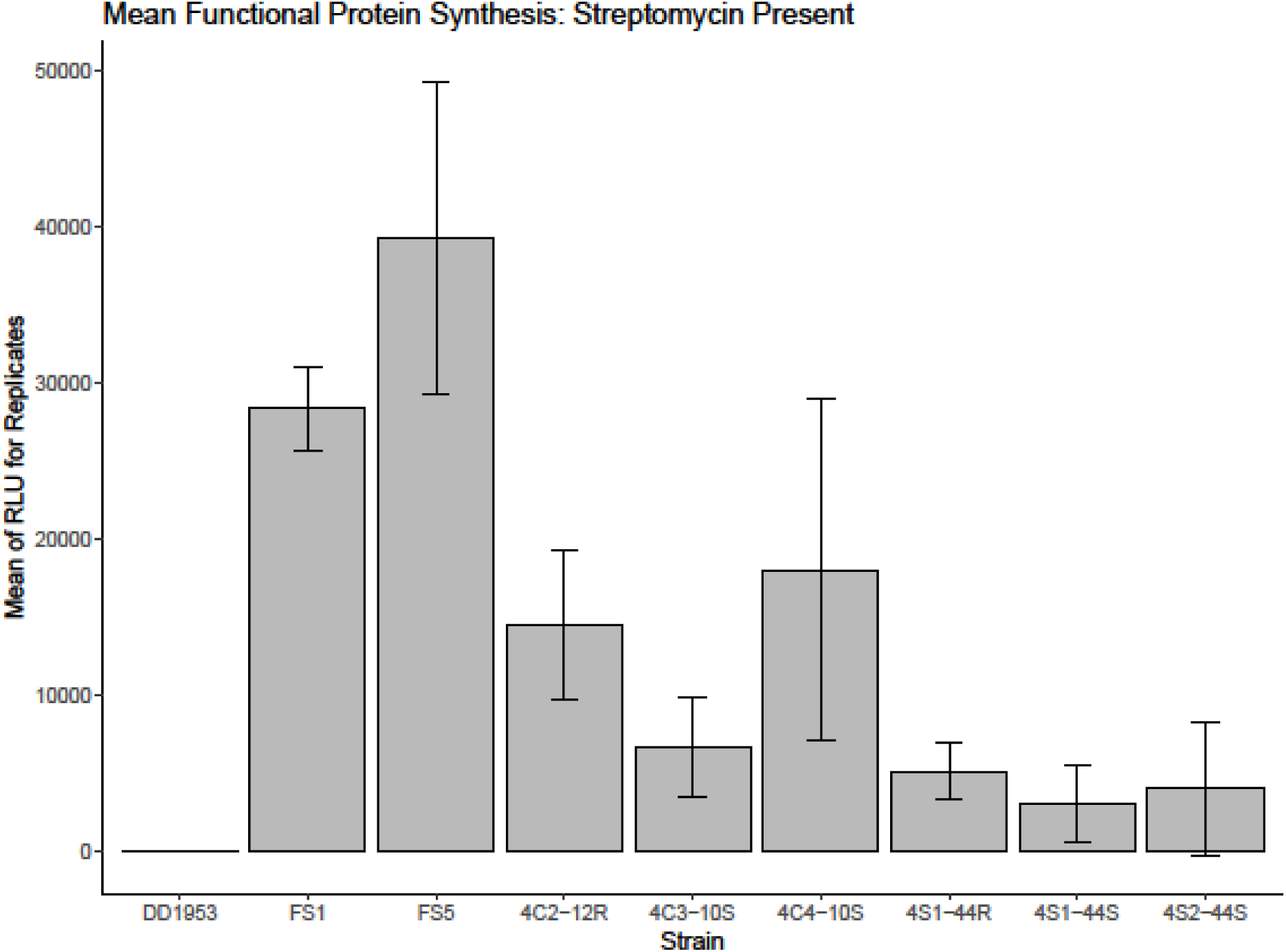
Mean Functional Protein Synthesis with Streptomycin Present. A functional protein synthesis assay was developed that quantified the amount of functional firefly luciferase protein produced in one hour. Quantities taken using Relative Light Units in a luminometer. Contrary to expectation, evolved strains had lower rates of function protein production than streptomycin resistant ancestors, FS1 or FS5. The evolved strain with the highest mean functional protein production (and largest confidence interval) is 4C4-10S, which evolved *without* streptomycin present in the chemostat. Error bars indicate 95% CI. See Supplemental Information for assay conditions and protocol.

## Discussion

Based upon previously published work, many of our results were precisely as expected for an experimental evolution study investigating adaptation post-resistance. We observed the cost of resistance typical of streptomycin resistance in *E. coli* with subsequent increases in relative fitness. In flasks, these fitness increases came via compensatory evolution at known loci. In chemostat evolved strains, fitness increased through changes in glucose uptake systems, which has been previously observed for *E. coli* evolved under glucose limited conditions, although not in relation to streptomycin adaptation. However, when we sought to investigate the mechanisms underlying these fitness increases, we found novel results suggesting that the ways in which antibiotic resistant strains adapt post-resistance is more complex than what is expected. We attempt to dissect our results and provide some potential context here.

General changes in fitness were anticipated: fitness relative to the antibiotic resistant ancestor (FS1, FS5) increased across all experimental populations. We found a strong signal for directional selection for increased time spent in exponential growth in all evolved populations even though the mechanism of this selection differs depending upon the environmental conditions experienced during the evolution experiment. This phenotype is one we do not believe has been highlighted before, but may play a large role in how bacterial populations grow, particularly if they are also antibiotic resistant. The environmental differences between conditions may seem minor, given that they were identical except for the *in vitro* method used and the streptomycin concentration (for chemostats), but even at this extremely fine scale, environment made all the difference.

In chemostats, the genotypic changes centered on glucose uptake involving the *galS* system. Similar changes have been observed previously in *E*.*coli* growing under glucose limited conditions in chemostats (Notley-McRobb and Ferenci 1999), however, in that case the strain of *E. coli* used, BW2952 (Genbank Accession CP001396), was not resistant to streptomycin. This tells us that such glucose uptake changes mean that this evolutionary path is accessible via selection without regard to antibiotic resistance phenotypes and was a result of the glucose limited environment.

In flasks, however, the mechanism was completely different. Genetic changes in serial transfer populations centered on additional changes to the strep-resistant ribosome. Similar compensatory mutations have been previously reported for streptomycin resistant strains of both *E. coli* (Schrag and Perrot 1996; Schrag et al. 1997) and *Salmonella Typhimurium* (Björkman et al. 1999, 2000). In the previous cases, these compensatory mutations have been shown to increase rates of polypeptide chain synthesis in the ribosome (Andersson et al. 1982), which is the phenotype most notably affected by the molecular mechanism of streptomycin resistance in those ribosomes. The speed of protein synthesis is integral to bacterial growth, and therefore fitness, in flasks and bacteria are expected to select for increases in maximum growth rate (μ_max_) under these conditions (Lenski 2017). Compensatory mutations in strep-resistant strains is thought to aid in increasing μ_max_.

However, when we directly measure μ_max_ in our flask evolved strains, the data do not support this hypothesis. FS1 and FS5 do not have lower μ_max_ values than the streptomycin sensitive ancestor, DD1953, even though they have a mutation in the *rpsL* gene known to impact protein synthesis. Additionally, flask evolved strains do not show increases in *µ*_max_ values, even though these populations have previously identified compensatory mutations (Table 2). In fact, there is no pattern of increase in *µ*_max_ for any of our experimentally evolved strains (Figure 1B). The lack of change in *µ*_max_ values is also supported by our functional protein synthesis assay, which unexpectedly found *lower* rates of production of functional protein in evolved strains than in the streptomycin resistant ancestor for all strains evolved in the presence of streptomycin (Figures 2, S1). In fact, the only evolved strain that did not have a significantly lower rate of protein synthesis was 4C4-10S, which was evolved without streptomycin *in a chemostat*.

Across all of our evolved strains, the only pattern of selection was for increases in the amount of time spent in exponential growth (Figure 1D). This signal was robust to all experimental conditions and concentrations of streptomycin. At the endpoints of the experiments, the time spent in exponential growth for evolved strains was similar to that for DD1953, suggesting that this phenotype has a large impact on relative fitness, a conclusion supported by our fitness assays.

In order to understand our results in context, we believe it is necessary to consider both the evolvability (Wagner and Altenberg 1996) as well as the fitness landscapes (Payne and Wagner 2019) of our strains in the environments tested. In chemostats, the genetic changes (*galS*) were predictable based upon the experimental conditions, even if the phenotypic changes (time in exponential growth) were not. Genotypic changes did not depend on the antibiotic resistance state of the genetic background and reflected adaptation to the limiting nutrient in a chemostat (Warsi and Dykhuizen 2017). In flasks, the selection was for genotypic changes that were not accessible via natural selection prior to streptomycin resistance. These mutations were, however, open to selection *after* the switch to a streptomycin laden environment, suggesting that the antibiotic resistance mutation not only allowed for the strain to survive in the new environment, but it also increased the evolvability of the strain in this new environment by opening up adaptive paths that did not exist beforehand. In doing so, post-resistance adaptation potential (i.e., the number of possible adaptive paths accessible by selection) increased. This makes the success of any anti-selection regime, such as those often relied upon in order to control the spread of resistance, highly improbable.

Considered in a different way, we understand that the addition of antibiotic changes the environment substantially, which also changes the adaptive landscape from the perspective of the bacteria. Strains that were high fitness in the strep-free environment cannot survive after the addition of streptomycin. This may be most evident in a genotype to fitness map, where we understand that the antibiotic sensitive ancestor, DD1953, was high fitness in the absence of streptomycin. However, the DD1953 strain has an effective fitness of 0 once streptomycin is added to the environment. In strep-laden environments, FS1 and FS5, which are lower fitness in the antibiotic free environment, assume higher fitness. The environmental change further reveals hidden genetic variation (Szamecz et al. 2014), offering FS1 and FS5 access to higher fitness optima via selection. Our data show that these fitness peaks are related to the time spent in exponential growth phenotype. Importantly, our results indicate that *how* strains access these peaks differs based upon their specific *in vitro* environments, with chemostats having an adaptive path similar to that available to strep-sensitive populations and batch cultures having an adaptive path contingent upon the strep-resistance mechanism, which was not available to strep-sensitive strains. In this way, phenotype and environment conspired to increase evolvability in our streptomycin resistant strains.

The increase in the number of possible adaptive paths for antibiotic resistant strains may provide a hypothesis to explain why selection against antibiotic resistance has proven so difficult. Populations at global optima, such as antibiotic sensitive strains, are likely to have inherent phenotypic compromises. For a large population with sizeable genetic diversity, environmentally mediated release from a global optimum can be expected to allow for directional selection for any variety of previously inaccessible phenotypes. Such a shift and change in accessibility reflects an increase in variability (Payne and Wagner 2019). Antibiotic resistance represents a specialized case of such an environmentally mediated release whereby the effect upon selection is particularly strong, with high fitness genotypes suddenly becoming lethal.

If evolvability does increase post-resistance, antibiotic rich environments may act as a mechanism that releases populations from the generalized fitness peaks upon which they had been stranded and may further provide additional paths with which they could then explore a newly enriched fitness landscape. By changing allelic frequencies while also potentially simplifying rough adaptive landscapes, the environmental shift increases variability and, therefore, evolvability. The increased availability of high fitness peaks, particularly in new environments, illustrates what we have come to think of as a rabbit hole effect, whereby previously unavailable genotype/fitness combinations become available through a sudden change in the environment. Much like Alice after tumbling to the bottom of the rabbit hole, there are new worlds suddenly available for exploration. In this study, the higher fitness flask evolved strains reached their fitness peaks by increasing the time spent in exponential growth relative to their direct strep-resistant ancestors through additional changes to a mutated ribosome. These changes were not genetically possible through selection without the sequence of resistance mutation, streptomycin positive environment, and then adaptation in batch culture. When the identical sequence is repeated in continuous culture, such genotypic changes are not observed, suggesting such a path is less likely, if not altogether impossible.

In 1975, H.W. Smith warned against the idea of combating established antibiotic resistance by expecting resistant strains to select for susceptibility simply by removing the antibiotic from the environment (Smith 1975). This ecological fallacy still remains and we have learned by some tough lessons that reversing resistance will not be quite as simple as hoped (Andersson and Hughes 2010). Recently, however, new approaches have begun to show promise in reversing resistance, such as in some phage treatments for drug resistant bacterial populations (Chan et al. 2016). This study suggests that undoing what has been done may be even more complicated yet: ephemeral switches in environments may increase the adaptive potential of resistant populations and allow for increases in fitness that cannot be predicted *a priori*.

## Acknowledgements

This work was supported by Lawrence B. Slobodkin and George C. Williams Fellowships (FS) and the Columbia Frontiers of Science Fellowship (FS). The authors thank Walt Eanes, John True, Peter Tonge, and Omar Warsi for helpful discussions and William Lannon and the Biology Department at Columbia for the generous use of department equipment. This is contribution number 1,247 from the Ecology and Evolution Department of Stony Brook University.

