## Supplemental Information for "Antibiotic Resistance Increases Evolvability and Maximizes Opportunities Across Fitness Landscapes"

For

Methods

**Strain Nomenclature**

Evolved strains are named according to the method of their experimental evolution vessel, the number of days that population was evolved, and their bacteriophage T5 resistance phenotype. For instance, strain 4C2-12R is the strain evolved in experiment series *4* in *C*hemostat *2* and was sampled on day *12* of the experiment and is bacteriophage T5 *R*esistant. In the case of flask evolved strains, the nomenclature is consistent, with the chemostat “C” replaced with an “S” for *S*erial transfer.

**Chemostats**

All continuous culture experiments were conducted in Kubitschek style glass chemostats (see Dykhuizen 1993 for in depth description). Chemostats constantly supply a set amount of fresh media to the 30 mL growth chamber. The amount of media supplied per unit time (the dilution rate, D) was controlled using a Wizco peristaltic pump. By controlling the amount of media available, the population growth rate was maintained in exponential phase throughout the experiment and the generation time was controlled. Here, the generation time was set to 3.69 hours per generation (6.5 generations per day, D = 0.1875), which matched the number of generations per day for flask evolved cultures. The media for all experiments was Davis salts supplemented with 0.01% glucose (w/v) as the sole available carbon source. Used media and dead cells left the growth chamber as fresh media entered. Removal was by means of a waste port on the side of the chemostat. All media and growing cells in the chemostat were constantly mixed via purified and humidified air pumped through the chemostat by an aquarium air pump so as to maintain proper mixing and to inhibit formation of biofilm on growth chamber walls. Chemostats were maintained at a constant 37°C temperature by means of a water bath.

Chemostats were initially set up and allowed to run for at least 36 hours prior to inoculation with the experimental populations. A sample was taken from the chemostats just before inoculation and plated, without dilution, to verify that the chemostats were free from contamination at the start. The inoculating sample was grown from frozen stocks overnight in flasks with media identical to that in the chemostat except for a 10-fold increase in the glucose concentration. To inoculate, 100 μL of the overnight growth was placed into 10 mL of fresh media with 0.1% glucose in a sterile side arm flask. Growth was tracked by measuring optical density at 600 nm with a WPA biowave model CO8000 Cell Density Meter. When the OD_600_ measured 1.7-1.85, the samples were immediately used to inoculate the chemostats. One mL of each strain of this inoculant was used in each chemostat.

Experimental chemostats were maintained until one strain reached either fixation or high-frequency. Samples were collected from the chemostats via the sampling port approximately every 24 hours starting from the time of inoculation. The chemostat samples were diluted 100,000 fold in Davis salts and plated in quadruplicate on LB agar plates with streptomycin concentrations matching the experimental condition. When bacteriophageT5 was being used as a molecular marker, the samples were also quadruplicate plated on LB agar plates supplemented with streptomycin and T5 bacteriophage (MOI>>2). All plates were incubated overnight at 37°C and CFUs were counted using a ProtoCol automatic plate reader and ProtoCol software V3.15, both by Synbiosis (Cambridge, UK). This procedure followed the method previously described (Dykhuizen and Hartl 1981). Basically, plates were made in quadruplicate and the values used in tracking the populations are the average of the 4 replicate plates. T5R and T5S counts were calculated by subtracting the mean number of T5R CFUs from the mean total of all CFUs.

Long term experiment population frequencies were tracked over the course of the experiments. As adaptive changes were selected for in the chemostat, the frequency of the clone with a positive adaptive mutation increased in frequency until a selective sweep was completed or another positive mutation came into a competing clone and the difference in average fitness between the two competing clones was lowered, in which case, the frequency of the second positive allele would rise in the population. If a second positive mutation occurred in the same clone as the first positive mutation, the rate of change in that clone would increase at the expense of other competing clones in the chemostat since the chemostat has a constant and limited population size. For *E. coli*, that population size in 30 mL chemostats with the media used here is 10^8^ cells/mL (N = 3.0 x 10^9^).

Chemostats 4C1 (data not reported here) & 4C2 utilized the Davis minimal media supplemented with 0.01% glucose and contained a concentration of 100 μg/mL of streptomycin in the media. Chemostat 4C3 had the Davis minimal media, 0.01% glucose, and 16 μg/mL of streptomycin. This concentration of streptomycin was chosen in part based upon the expected MIC of *E. coli* as per Miller (Miller 1992). Chemostat 4C4 served as a negative control: Davis minimal media with glucose, but no streptomycin.

Chemostats were changed and the experimental media and population was moved into new sterile chemostats every 10 days or less, so as to avoid the complicating factors associated with any possible biofilm formation on chemostat walls. Media and media jars were also refreshed every week. All streptomycin came from a single batch of prepared stock (10 mg/mL).

**S30 Lysate Preparation**

This protocol is adapted (Shrestha et al. 2012) and uses a cup horn sonicator to prepare high quality *E. coli* cell extracts. The protocol can also be found on corresponding author’s website:

https://fabspagnolo.wordpress.com/resources/

Grow *E. coli* cells overnight in 3 mL of 2XYT broth with MOPS,

Pipet 0.5 mL of overnight growth into 12 mL of fresh 2XYT with MOPS,

Take optical density reading of your fresh sample and zero out your OD reader to standardize on this sample,

Incubate at 37° C (constant stirring),

Ice your sample,

When the OD reaches 0.8, take a sample for plating (to quantify number of cells prior to sonication),

Place the 12 mL of growth into 12 1.5 mL centrifuge tubes

Centrifuge (at 4° C) for 15 minutes at 14,000 RPM,

Remove supernatant,

Wash pellet with 1 mL of ice cold S30 Lysate Buffer A ,

Vortex to resuspend pellet,

Centrifuge again for 15 minutes at 14,000 RPM,

Repeat steps above for a second wash,

After 2^nd^ wash, consolidate pellets by placing 3 pellets into a single 1.5 mL centrifuge tube and resuspend in 1 mL of fresh S30 Lysate Buffer A (ice cold) so that you now have 4 centrifuge tubes per sample rather than 12,

Flash freeze in liquid nitrogen. If you desire, you can store these frozen samples overnight at -80° C and continue with sonication step following day,

When you are ready for sonication using a cup horn sonicator (a high capacity chiller is preferred as sonication medium will increase in temperature; be sure to monitor temperatures), thaw your samples and place in sonication chamber of chilled (4° C) cup horn sonicator,

Sonicate at 75% amplitude, 30 seconds on and then 30 seconds off for a total of 40 minutes of sonication (this is 40 minutes of total “on” time). Monitor the temperature of the samples. If temperature climbs too quickly, at the 20 minute mark, rest samples and allow the chiller to cool the samples down,

Plate a sample post sonication to quantify the efficiency of sonication. Yield can be improved by adjusting total sonication time, if needed.

Post sonication samples can be stored at -80° C.

*2XYT with MOPS media:*

Start with DDH_2_0. For 1 liter of broth, add:

16 grams Tryptone

10 grams Yeast Extract

5 grams NaCl

23 grams of MOPS (FW 231.25)

*S30 Lysis Buffer A:*

Start with DDH_2_0. For 1 liter of buffer, add:

1.2 grams TRIS Base (FW 121.1, for 10 mM)

2.6 grams Magnesium Acetate (FW 214.45, for 12 mM)

60 mL of 1M Potassium Glutamate (for 60 mM)

0.15 grams DTT (FW 154.25, for 1 mM)

**Protein Synthesis Assay Protocol**

The protein synthesis assay protocol is adapted from the Promega protocols for the *E. coli* S30 Extract System for Circular DNA (firefly luciferase control reaction) and the Promega Bright-Glo Luciferase Assay System. This assay is based on methods previously used for ribosomal polypeptide synthesis assays (such as in Holberger and Hayes, 2009). The complete protocol can also be found on corresponding author’s website:

https://fabspagnolo.wordpress.com/resources/

From the Promega *E. coli* S30 Extract System for Circular DNA kit (Cat. No. L1020),

Mix the 3 partial Amino Acid mixtures (from kit) together for a total AA mix (all at [1 mM]) Total vol = 525 ul

Mix the 2 containers of pBESTLuc plasmids (10 ul of conc 1 ug/ul each) then add 60 ul H_2_0 for total vol of 80 ul (final concentration is 0.25 ug/ul)

In order to make Master Mix, add:

200 ul	Total AA Mix [1 mM]

750 ul	S30 Premix without AAs (contains: tRNAs, ATP, rNTPs)(from kit)

50 ul	molecular grade H_2_O

*Reaction Mix (each 1.5 mL tube):*

35 ul	Master Mix

1 ul	pBESTLuc Plasmid prep

60 ul	your S30 Lysate sample

5 ul Molecular grade Water (OR, if you are performing the assay in presence of antibiotic such as streptomycin: 5 ul of [2 ug/uL] Streptomycin)

Make 3 replicates of each S30 lysate sample/reaction mix,

Add plasmid last and immediately, vortex,

Incubate at 37 C for 60 min,

Ice at exactly 60 minutes,

Add 25 ul of Bright Glo Luciferase Assay Reagent, make sure that reagent is at room temperature,

Place 1.5 mL tube into Luminometer,

Read Raw Data Measurement in RLU (3 times for each sample is recommended).

Positive controls can be performed by using the prepared S30 Lysate from the kit as a substitute for your S30 lysate sample.

Negative controls should be run under a range of conditions that allow you to verify that you will not have positive RLU reads from any combination of the reagents used, including the H_2_O.

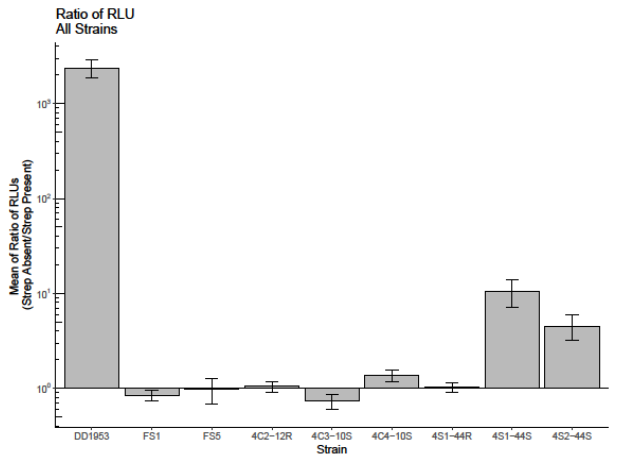

**Figure S1: Ratio of Functional Protein Synthesis in Absence/Presence of Streptomycin**

The protein synthesis assay was repeated in both the absence and presence of streptomycin in order to quantify the effect of the antibiotic on protein synthesis *in vitro*. For DD1953, the large effect is to be expected. DD1953 is streptomycin sensitive and WT ribosomes should not be able to produce proteins when streptomycin is present. For most of the evolved strains, the ratio of relative light units produced from functional luciferase is approximately 1:1. The exceptions to this are both flask evolved (4S1-44S and 4S2-44S). Both of these strains produced significantly more functional protein in the absence of streptomycin than when it was present. It should be noted, however, that these strains evolved in the presence of streptomycin for almost 300 generations, suggesting that the observed effect is pleiotropic and not selected. Error bars represent 95% Confidence Intervals.
